# Correlating single-molecule characteristics of the yeast aquaglyceroporin Fps1 with environmental perturbations directly in living cells

**DOI:** 10.1101/2020.02.27.967638

**Authors:** Sviatlana Shashkova, Mikael Andersson, Stefan Hohmann, Mark C Leake

## Abstract

Membrane proteins play key roles at the interface between the cell and its environment by mediating selective import and export of molecules via plasma membrane channels. Despite a multitude of studies on transmembrane channels, understanding of their dynamics directly within living systems is limited. To address this, we correlated molecular scale information from living cells with real time changes to their microenvironment. We employed super-resolved millisecond fluorescence microscopy with a single-molecule sensitivity, to track labelled molecules of interest in real time. We use as example the aquaglyceroporin Fps1 in the yeast *Saccharomyces cerevisiae* to dissect and correlate its stoichiometry and molecular turnover kinetics with various extracellular conditions. In this way we shed new light on aspects of architecture and dynamics of glycerol-permeable plasma membrane channels.

## Introduction

Traditional techniques in biochemistry and molecular biology are usually performed on a population ensemble average level. Such approaches “smooth” the noise by averaging the observations from anomalous outlying units. Every population under investigation, whether it is a cell culture or a protein bulk within a unicellular organism, is a heterogeneous system. Therefore, ensemble averaging masks important effects of subpopulations ^1–3^, such as drug resistant bacteria or cancer cells ^4–6^. Single-molecule optical biophysics methods permit real-time visualization of key cellular processes ^7^, the action of so-called biological ‘nanomachines’ ^8^, such as signal transduction, gene expression, immune response, mapping cellular genome, etc., providing direct insights into molecular mobility, stoichiometry, copy numbers ^9,10^. Novel localization-based super-resolved microscopy techniques allow tracking individual molecules of the same type (e.g. FliM protein of *Escherichia coli* ^8^) or different types (e.g. correlating separate motions of proteins and lipids in the same bacteria cell ^11^) to uncover “hidden” subpopulations, thus, determining precise biological functions ^9^.

An important part in understanding the regulation of a membrane protein is revealing its dynamics of within the membrane, interaction with other proteins, re-localization and turnover ^9^. One of the most frequently used super-resolution methods for plasma membrane protein studies is total internal reflection fluorescence (TIRF) due to its relative simplicity to setup to enable selective illumination and excitation of fluorophores positioned close to the cover slip. Photoactivated localization microscopy (PALM) and stochastic optical reconstruction microscopy (STORM) have a wide range of applications including membrane protein investigations too ^4^. Conventional PALM and STORM utilize a photoconversion process of the fluorophore and involve reconstruction over normally thousands of consecutive image frames. These techniques have a typical effective temporal resolution of 0.5-1 s which provides limitations in studying fast dynamic processes in living cells ^12,13^. However, more rapid sptPALM approaches have recently been employed to monitor dynamics with a time resolution of a few tens of ms per frame ^14^.

The plasma membrane is a selectively-permeable natural barrier that ensures cell homeostasis via controlling among others nutrient sensing and uptake as well as flux of metabolites ^15–17^. The Fps1 protein of the budding yeast *Saccharomyces cerevisiae* is a gated aquaglyceroporin, a member of the major intrinsic protein (MIP) family of plasma membrane channel proteins. The primary purpose of Fps1 is to mediate the efflux of glycerol to regulate cellular turgor pressure ^18^. Fps1 is actively regulated in response to hyper and hypo-osmotic stress, or presence of compounds in the environment to which Fps1 is permeable ^18–21^.

Studies on Fps1 channel stoichiometry via co-immunoprecipitation followed by SDS-PAGE separation and immunoblot analysis suggest that Fps1 exists as a monomer but may also self-associate into multimeric complexes with up to four Fps1 molecules ^22^. Structural analyses on other MIP members, canonically water permeable aquaporins ^23^, indicate a tetrameric organization where each monomer defines a pore with a possible fifth pore formed in the center of the tetramer ^24^. However, the aspects of MIP channels assembly and architecture, dependencies of their regulation in living cells on the microenvironment, remain to be elucidated.

Here, we employ single-molecule Slimfield super-resolved fluorescence microscopy on GFP-tagged yeast Fps1 to reveal new aspects of the MIP proteins organization and regulation which links directly to their function and, importantly, correlate our observations to changes in the extracellular microenvironment. Slimfield is a powerful optical microscopy tool that uses spatially delimited illumination confined to vicinity of approximately a single cell and enables millisecond imaging of fluorescent protein fusions directly in living cells ^25–27^. We have also combined this technique with deconvolution analysis to calculate Fps1 copy numbers. Our study provides novel insights into understanding of cellular adaptation to the microenvironment through characterization of the plasma membrane channels.

## Material and methods

### Growth conditions and media

Cells from frozen stocks were pre-grown on standard YPD medium (20 g/L Bacto Peptone, 10 g/L Yeast Extract) supplemented with 2% glucose (w/v) at 30°C overnight. For liquid cultures, cells were grown in Yeast Nitrogen Base (YNB) medium (1x Difco™ YNB base, 1x Formedium™ complete amino acid Supplement Mixture, 5.0 g/L ammonium sulfate, pH 5.8-6.0) supplemented with 2% glucose (w/v) and 1M sorbitol if required, at 30°C, 180rpm.

For microscopy experiments, cells were pre-grown overnight in YNB media with 20 g/L glucose and grown until mid-logarithmic phase, OD_600_ 0.4-0.7. For sorbitol experiments, all media were supplemented with 1M sorbitol throughout the whole process to reach full osmoadaptation to the high osmolarity. For oxidative stress experiments, cultures were treated with 0.4mM H_2_O_2_ immediately prior to imaging. Cells were then immobilized by placing 5µL of the cell culture onto a 1% agarose pad perfused with YNB supplemented with 2% glucose (w/v) and 1M sorbitol or 0.4mM H_2_O_2_. The pad with cells was sealed with a plasma-cleaned BK7 glass microscope coverslip (22×50 mm).

### Strain construction

To prevent artificial aggregation of GFP, we used a variant (further denoted as mGFP) containing a A206K mutation to discourage self-oligomerization, as well as S65T and S72A mutations to improve protein photostability and fluorescence output ^4,28,29^. Although within protein fusions, large fluorescent proteins may disturb physiological behavior of a protein under investigation, functionality of fluorescently labelled Fps1 has been previously confirmed ^19,30–32^.

mGFP-HIS3 fragment from pmGFP-S plasmid ^33^, flanked with 50bp up- and downstream the STOP codon of the *FPS1* gene on 5’ and 3’ ends, respectively, was amplified by PCR using 200 µM of each dNTP, forward and reverse primers 0.5 µM each, 1x Phusion HF buffer (Thermo Scientific), 0.02 U/µL Phusion™ High-Fidelity DNA Polymerase (Thermo Scientific) reaction mix (for PCR program details see Supplementary Table 1) and purified with QIAquick PCR Purification Kit (QIAGEN). The Fps1-mGFP strain was created by transforming the BY4741 background strain with 100 µL of the purified mGFP-HIS3 fragment with standard LiAc protocol to allow for homologous recombination ^34^. Successful clones were verified by the confirmation PCR and standard epifluorescence microscopy. PCR primers used in this study are listed in Supplementary Table 2.

### Single-molecule Slimfield microscopy

Slimfield excitation was implemented via 50mW 473 nm wavelength laser (Vortran Laser Technology, Inc.) de-expanded to direct a beam onto the sample at 15mW excitation intensity to observe single GFP in living yeast cells ^33^. Fluorescence emission was captured by a 1.49 NA oil immersion objective lens (Nikon) followed by 300 mm focal length tube lens (Thorlabs) ^10^. Images were collected at 5ms exposure time every 10ms by Photometrics Evolve 512 Delta EMCCD camera using 93 nm/pixel magnification.

The focal plane was set to mid-cell height using the brightfield appearance of cells. As photobleaching of mGFP proceeded during Slimfield excitation distinct fluorescent foci could be observed of half width at half maximum 250-300 nm, consistent with the diffraction-limited point spread function of our microscope system, which were tracked and characterized in terms of their stoichiometry and apparent microscopic diffusion coefficient. Distinct fluorescent foci that were detected within the microscope’s depth of field could be tracked for up to several hundred ms, to a super-resolved lateral precision ∼40 nm ^35^ using a bespoke single particle tracking software written in MATLAB (MATHWORKS) and adapted from similar live cell single-molecule studies ^10,35–37^.

The molecular stoichiometry for each track was determined by dividing the summed pixel intensity values associated with the initial unbleached brightness of each foci by the brightness corresponding to that calculated for a single fluorescent protein molecule (mGFP for 473 nm wavelength excitation) measured using a step-wise photobleaching technique described elsewhere ^33,38^. The apparent microscopic diffusion coefficient *D* was determined for each track by calculating the initial gradient of the relation between the mean square displacement with respect to tracking time interval using the first 10 time intervals values while constraining the linear fit to pass through 4σ^2^ on the vertical axis corresponding to a time interval value of zero. Maturation effects of fluorescent protein fusions withing living cells were characterized on similar yeast cell lines previously, indicating typically 10-15% immature ‘dark’ fluorescent protein ^39^.

Similar to molecular stoichiometry calculations, total numbers of Fps1-mGFP were estimated based on the background- and autofluorescence corrected integrated density values of each cell as well as fluorescence intensity of a single fluorophore obtained through ImageJ Fiji Software.

## Results and discussion

### Fps1 reside in multi tetrameric assemblies

In both eukaryotes and prokaryotes, aquaglyceroporins have been reported to act as tetramers ^40–43^. To visualize and further investigate previous reports on Fps1 tetramerization ^22^, we employed single-colored single-molecule fluorescence Slimfield super-resolution microscopy (Fig. 1A) to determine the stoichiometry of the GFP-tagged yeast Fps1 under non-stressed conditions (Fig. 1A, top panel, Supplementary Video 1). Consistent with previously published data and observations of aquaporins in other eukaryotes, Fps1 seems to be present as tetramers, which are also organized in higher stoichiometry spots (Fig. 1C). Based on the previous estimation of the *S. cerevisiae* plasma membrane width ^44^, we accepted any GFP tracks found between the cell boundaries identified from the bright field image and *ca.* 7 nm into the cell as the plasma membrane foci, henceforth referred to as membrane, while the spots found in the rest of the cell as “intracellular”. The mean apparent stoichiometry of the membrane foci is higher compared to those found in the rest of the cell (14±6.2 and 12±6.3, respectively. Student *t*-test. Fig. 1D). Unlike membrane Fps1, intracellular spots seem to be also present as lower stoichiometry oligomers (Fig. 1D, inset, and 1E). Foci of the same stoichiometry can be both immobile (diffusion coefficient, *D*<0.3 µm^2^/s) and fast moving (*D*>0.8 µm^2^/s) (Fig. 1E). Other aquaporins are known to reside in intracellular storage vesicles or lipid rafts until they are required at the plasma membrane ^45–47^. The abundance of intracellular Fps1 suggests that it could also reside in such vesicles. Key glycerol metabolizing enzymes are located in subcellular compartments, such as mitochondria, peroxisomes and lipid droplets ^48–50^. Thus, intracellular Fps1 might be sitting on their membranes where it mediates the flux for a substrate.

**Figure 1.**
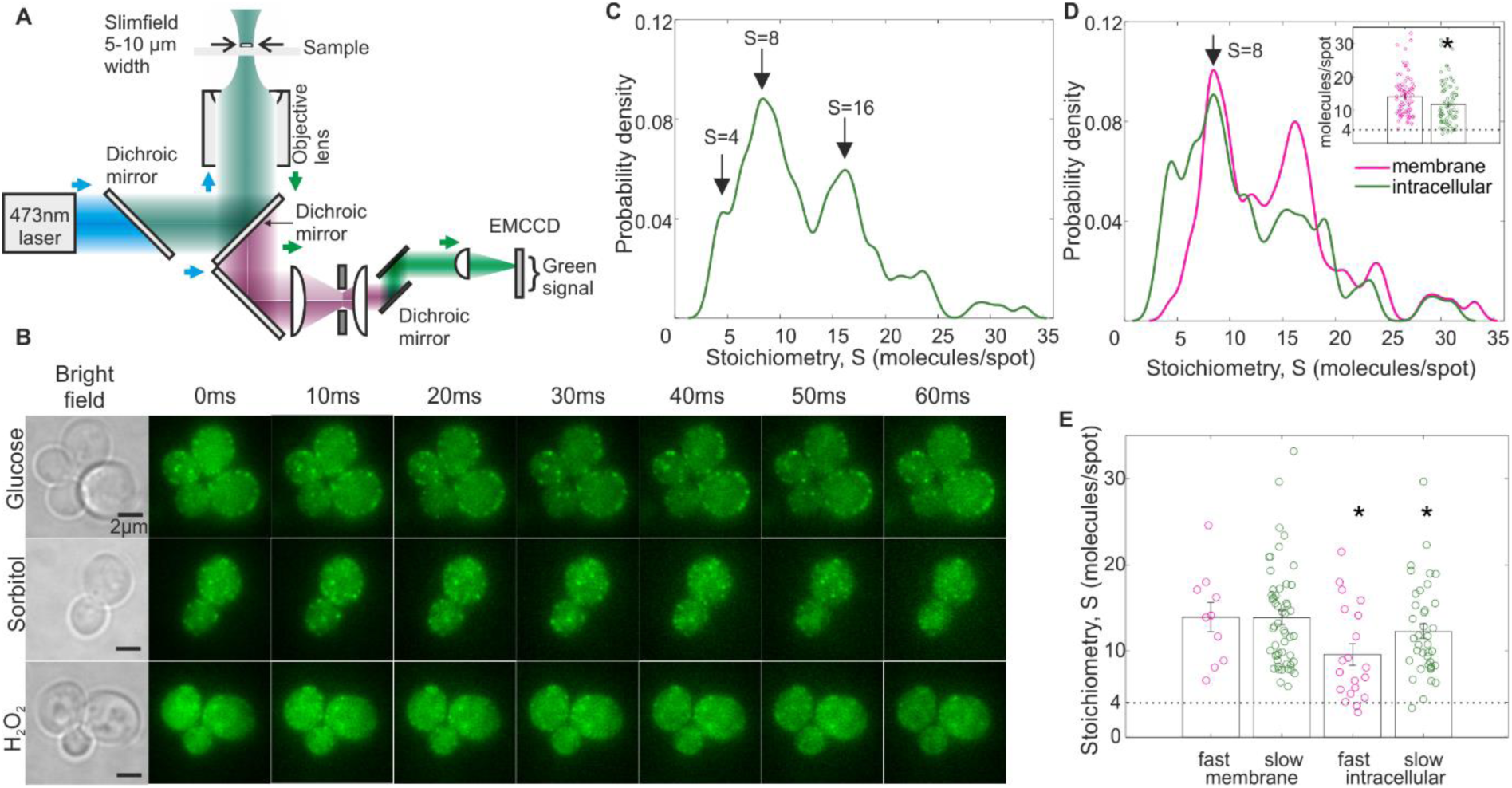
A. Single-colored Slimfield super-resolved microscopy setup. B. Examples of Slimfield images of S. cerevisiae cells expressing genomically integrated Fps1-mGFP fusion under non-stressed conditions (top panel, glucose), hyper-osmotic (middle panel, sorbitol) and oxidative stress (bottom panel, H_2_O_2_): bright field and green channel (right) are shown. Scale bar 2µm. C. Kernel Density Estimation plot (kernel width=0.7) of the Fps1-mGFP stoichiometry distribution under non-stressed conditions (n=40 cells). D. Kernel Density Estimation plots of the Fps1-mGFP stoichiometry distributions on the cellular membrane (magenta) and intracellular (green) spots. Inset: Jitter plots of membrane (magenta) and intracellular (green) Fps1-mGFP foci stoichiometry. Error bars represent standard error of mean. Student t-test, p<0.05. E. Jitter plots of apparent stoichiometries of fast moving (fast), diffusion coefficient, D>0.8 µm^2^/s, and immobile, D<0.3 µm^2^/s, (slow) membrane and intracellular Fps1-mGFP foci. Standard error bars are indicated. Student t-test, *p<0.05.

Interestingly, the mean apparent stoichiometry of the fast moving and immobile membrane foci is also similar (Fig. 1E). The limitation of our segmentation method is that budding cells are accepted as two separate cells with the “whole-cell” plasma membrane without accounting for the mother-daughter cell connection through the bud neck. Therefore, the mobile pool might represent Fps1 located closer to the bud, where proteins can move between the mother and the daughter cells, which would be consistent with previous reports on multiple plasma membrane proteins are asymmetrically segregated ^51^. However, further studies with bud neck labelling should be performed to verify that.

### Sorbitol causes a change in Fps1 organization

Biochemical studies suggest that increased external osmolarity causes rapid Fps1 closure, whereas decreased osmolarity results in the channel opening ^30,52,53^. To determine how hyper-osmotic conditions affect Fps1 architecture, we grew the cells in 1M sorbitol to reach complete osmoadaptation (Fig. 2A and Fig. 1B, middle panel, Supplementary Video 2). Sorbitol growth causes the apparent stoichiometry shift towards higher oligomeric clusters (Fig. 2B) with the highest probability peak of 8 molecules/spot in the absence of sorbitol (Fig. 1B) and 11 molecules/cell in sorbitol (Fig. 2C left). Sorbitol had been shown to increase molecular crowding in cells ^54,55^. The Fps1 protein contains 11 regions with intrinsic disorder making an overall proportion of disordered content >44%, more than 71% of which is within the large N- and C-terminal cytoplasmic domains (as predicted by PONDR software). Disordered motifs may undergo phase transition resulting in formation of higher oligomers, in this case, facilitated by increased intracellular crowding ^33^. These cytoplasmic domains are however also shown to be the main facilitators of protein interaction with Fps1 ^19,31,56–58^. Computer simulations indicate that specific protein interactions can also be stabilized upon an increase in molecular crowding ^59^. This together with known interactions with of Fps1 in membrane bound signal scaffolding is also likely to affect Fps1 stoichiometry ^56^. No clear periodicity of four Fps1 molecules/spot was found in sorbitol-grown cells. The overall stoichiometry spread is much broader compared to that found in glucose (Fig. 2B inset) regardless of the foci localization, indicating a range from Fps1 dimers up to 50 molecules/spot (Fig. 2C right).

**Figure 2.**
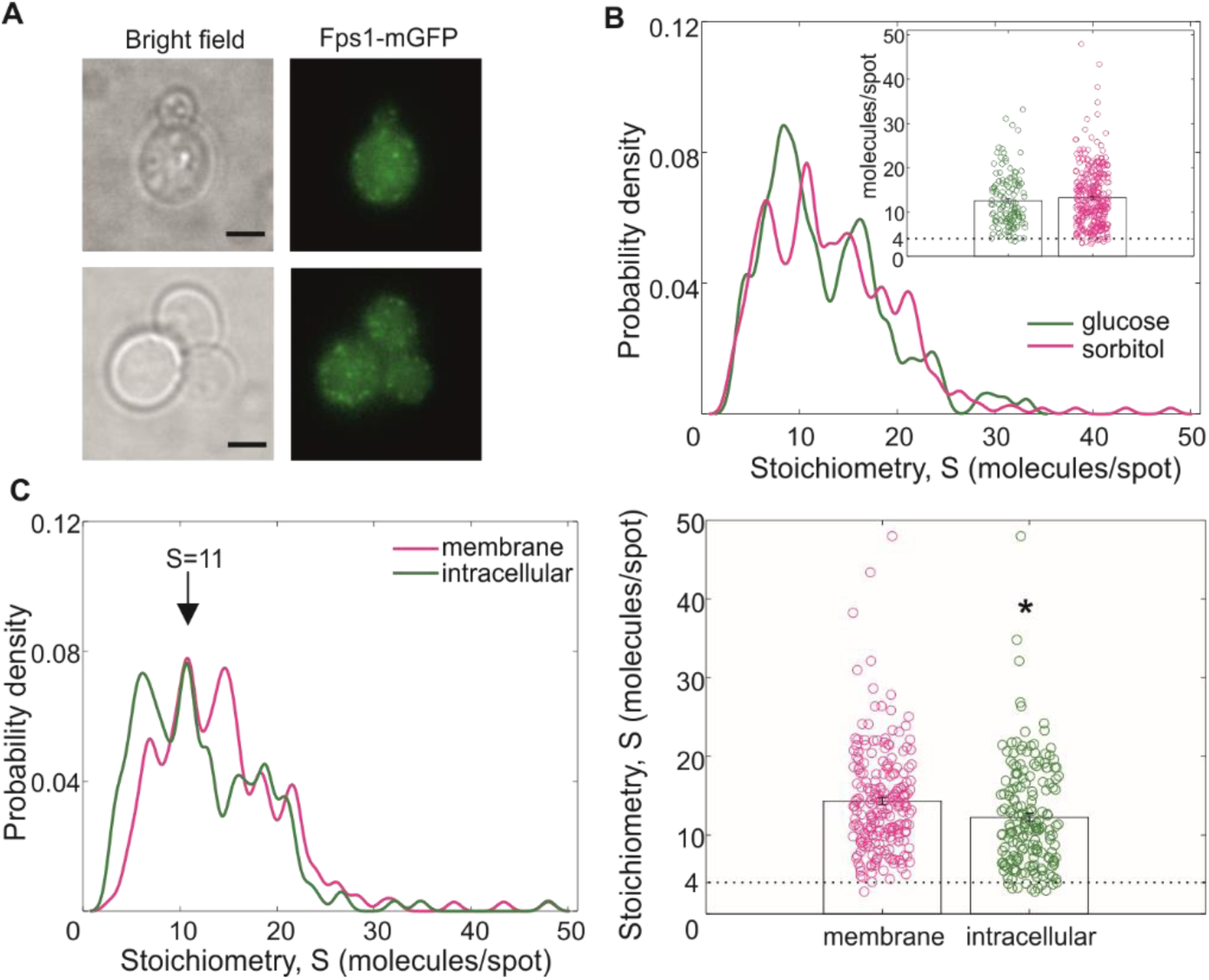
A. Examples of Slimfield images of S. cerevisiae cells expressing genomically integrated Fps1-mGFP fusion under hyper-osmotic stress (media supplemented with 1M Sorbitol): bright field (left) and green channel (right) are shown. Scale bar 2µm. B. Kernel Density Estimation plot (kernel width=0.7) of the Fps1-mGFP stoichiometry distributions upon normal (green, n=40 cells) and hyper-osmotic stress (magenta, n=89 cells) conditions. Inset: Jitter bar chart of mean apparent stoichiometry, standard error of mean error bars are shown. C. Left, Kernel Density Estimation plots of the Fps1-mGFP stoichiometry distributions on the cellular membrane (magenta) and intracellular (green) spots. Right, Jitter plot of membrane (magenta) and intracellular (green) Fps1-mGFP foci stoichiometry. Error bars represent standard error of mean. Student t-test, p<0.05.

Increased crowding also results in lower mean diffusion coefficient, *D*: 0.43±0.61 in non-stress conditions as opposed to 0.36±0.51 upon hyper-osmotic environment (Fig. 3A). However, no obvious linear correlation between the diffusion coefficient and foci stoichiometry was observed in normal and stress conditions (Fig. 3B). According with our expectation that molecular movement is more restricted in the membrane, only few of the “fast” spots (*D*>0.8 µm^2^/s) are found on the membrane under both non-stress (glucose) and sorbitol conditions (Fig. 3B top left and bottom left). “Immobile” foci (*D*<0.3 µm^2^/s) are equally present on the membrane and in the rest of the cell (Fig. 3B top right and bottom right). Interestingly, spots with the lowest D are not actually the ones of the highest stoichiometry. This might suggest that Fps1 is trafficked as, and thereby also likely regulated as, a multimer.

**Figure 3.**
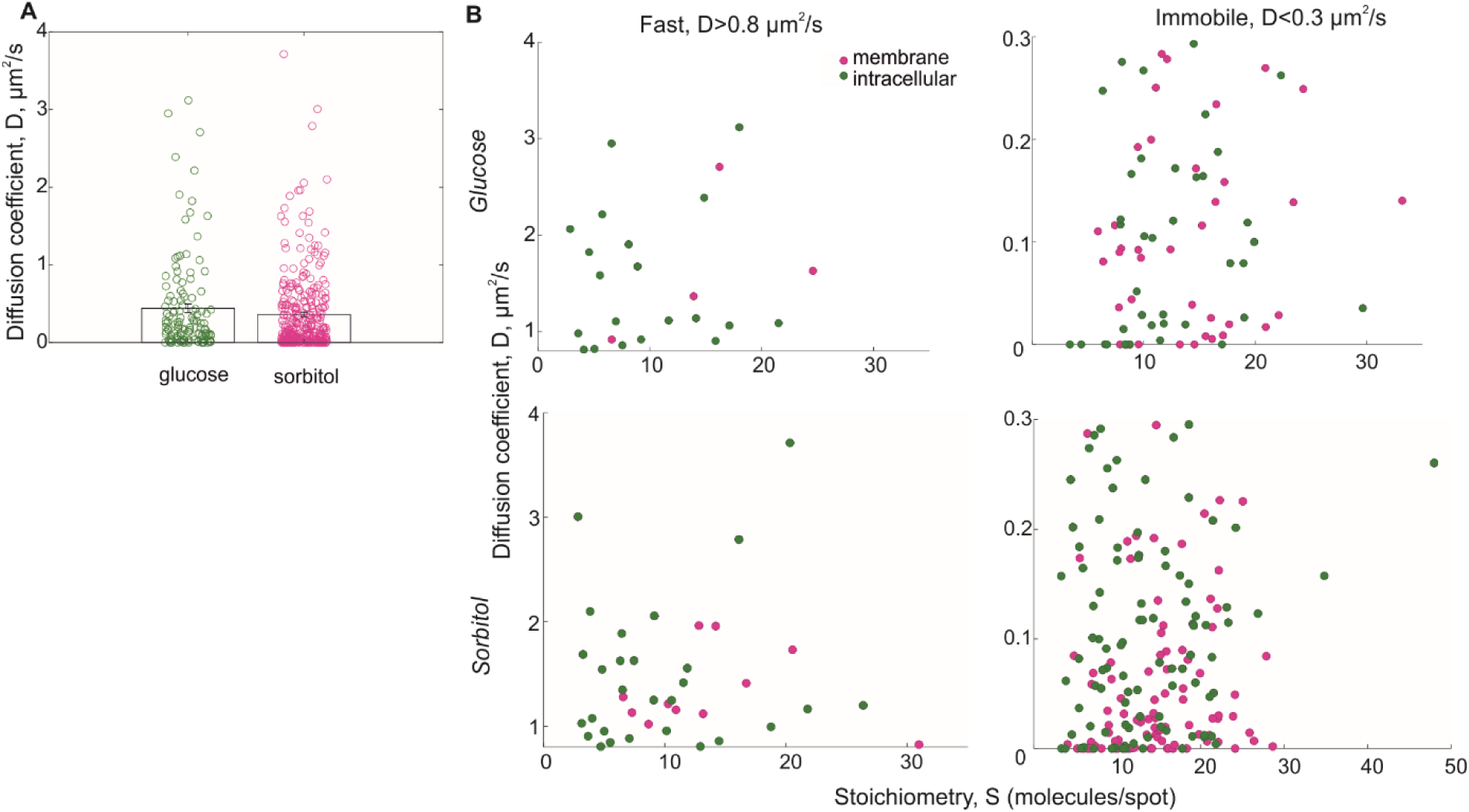
A. Jitter plot of the mean diffusion coefficients of Fps1-mGFP foci under normal (glucose, green) and hyper-osmotic stress (sorbitol, magenta) conditions. Error bars represent standard error of mean. B. Scatter plots of stoichiometry and diffusion of fast, D>0.8 µm^2^/s (left), and immobile, D<0.3 µm^2^/s (right), Fps1-mGFP spots found on the membrane (magenta) and the rest of the cell (green) upon non-stress (top) and hyper-osmotic (bottom) conditions.

### Oxidative stress rapidly facilitates Fps1 degradation

MIPs have been shown to facilitate hydrogen peroxide diffusion through cell membranes ^60–62^. To study the effect of the rapid exposure to the oxidative stress on Fps1 composition, we treated cells with 0.4mM H_2_O_2_ for 20-40 min prior imaging (Fig. 1B, bottom panel, Fig. 4A, Supplementary Video 3). No significant differences could be found in mean apparent stoichiometry and diffusion coefficients between all three conditions (Fig. 4B). Similar to that upon sorbitol treatment, regardless of the cellular compartment, Fps1 exists as a dimer (Fig. 4C).

**Figure 4.**
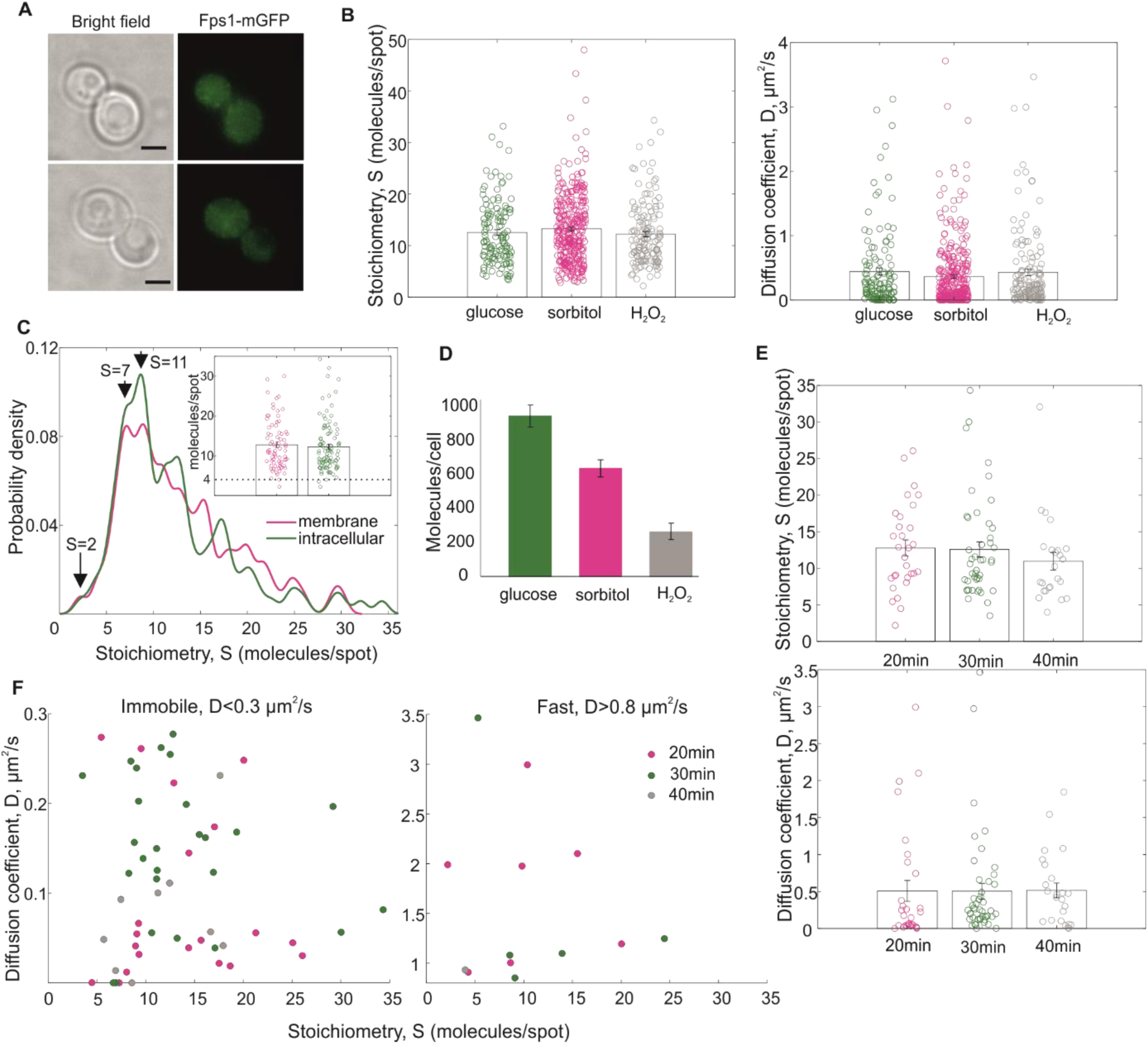
A. Examples of Slimfield images of S. cerevisiae cells expressing genomically integrated Fps1-mGFP fusion upon oxidative stress: bright field (left) and green channel (right) are shown. Scale bar 2µm. B. Jitter plots of Fps1-mGFP stoichiometry (left) and diffusion coefficient (right) under various conditions. Standard error bards are shown). C. Kernel Density plot of the Fps1-mGFP stoichiometry distributions on the cellular membrane (magenta) and intracellular (green) spots. Inset: Jitter bar chart of membrane (magenta) and intracellular (green) Fps1-mGFP foci stoichiometry. Error bars represent standard error of mean. D. Numbers of Fps1 molecules per cell in yeast grown in standard (glucose, green), osmotic (sorbitol, magenta) or peroxide stress (H_2_O_2_, grey) conditions. Standard error bars are indicated. E. Jitter plots of Fps1-mGFP foci stoichiometry (top) and diffusion coefficients (bottom) in cells treated with hydrogen peroxide for 20 min (magenta), 30 min (green) or 40 min (gray). Error bars represent standard error of mean. F. Scatter plots of immobile (left) and fast moving (right) Fps1 foci in cells exposed to H_2_O_2_ for 20 min (magenta), 30 min (green) or 40 min (grey).

We identified three times less Fps1 molecules in cells exposed to H_2_O_2_ compared to those incubated in non-stress conditions (Fig. 4D). Our result is consistent with previous reports on hydrogen peroxide inducing protein internalization or degradation in various organisms - in plants, aquaporins have been shown to internalize upon H_2_O_2_ and salt treatment ^46,62^, degradation has been demonstrated for hemoglobin and membrane proteins in mammals ^63^ and intracellular proteins in *E. coli* ^64^.

The presence of dimers during both sorbitol growth and H_2_O_2_ stress, together with lower copy numbers of Fps1 molecules (Fig. 4D), point towards differences in protein turnover during these conditions compared to the standard environment. Dimerization as an intermediate state has already been reported for MIPs in other organisms. For example, the tetramer of the *E.coli* aquaglyceroporin GlpF unfolds via a dimeric intermediate state ^65^. In plants, there is also evidence of constitutive aquaporin cycling which has been theorized to favor the plasticity of the channel activity and its response to sudden environmental changes ^46^. The dimerization of plant aquaporins is thought to stabilize the channel in the membrane ^66–69^. Thus, the different range of Fps1 stoichiometries under stress conditions indicates changes in protein turnover leading in the case of H_2_O_2_ to degradation.

To determine if the Fps1 spots’ behavior changes over time, we split the acquired dataset into three groups representing cells being exposed to the oxidative stress for 20, 30 or 40 min. Consistently with our expectations, the mean apparent stoichiometry of Fps1-mGFP seems to decrease over time (Fig. 4E top). Interestingly, the smallest stoichiometry number increases across all three groups, from two molecules/spot in group 1 (20min H_2_O_2_ treatment) to four in group 3 (40 min). While no difference in the mean diffusion coefficient was identified between three groups (Fig. 4E bottom), the number of fast moving (*D*>0.8µm^2^/s) Fps1 foci identified in group 3 cells (40 min hydrogen peroxide treatment) is significantly smaller (Fig. 4F). Together, these findings point towards the initial assembly of smaller foci into larger stoichiometry spots which are further subjected to hydrogen peroxide-induced degradation.

## Conclusions

In 1972 the idea that the cell membrane was a fluid two dimensional lipid bilayer was introduced ^70^. Since then our knowledge of the plasma membrane and its components has increased exponentially and we know that it is a complex and dynamic network of lipids and proteins that can cluster and confer general and localized properties in both two and three dimensions ^71,72^. These protein clusters can either be stable or exist more transiently ^73,74^ and facilitate various cellular processes, such as signal transduction and gene regulation via recruitment of protein complex components to target DNA and channel activators to the plasma membrane ^31,33^.

Here we show that the yeast aquaglyceroporin Fps1 on the membrane is organized into multimeric clusters of varying sizes. Our data show that the oligomerization state of Fps1 alters depending on external stimuli. This is consistent with previous studies showing that the *E.coli* aquaglyceroporin GlpF has different oligomerization states depending on salt concentration ^75^. Rapid light microscopy approaches, such as those employed by us here, also permit investigating regulatory features on a single-molecule level which is difficult to achieve with traditional biochemical methods or standard epifluorescence imaging techniques. The key importance is correlating these molecular scale observations with changes to the microenvironment of individual cells. In doing so, we provide evidence that hyper-osmotic conditions and oxidative stress change the rates of Fps1 turnover compared to standard non-stress conditions. Moreover, we suggest that one of the steps in Fps1 degradation upon oxidative stress is the assembly of smaller foci into larger clusters. We also show, that inside the cell, Fps1 exists in an immobile state which might indicate its presence on membranes of intracellular organelles.

Our study provides novel insights into real-time MIPs dynamics within plasma membrane and in the rest of the cell. Further multicolor super-resolution millisecond approaches can be applied to the studied system to determine the machinery of channel interactions with associated proteins which regulate channel opening and closure or provide a binding scaffold, such as Sho1 protein scaffold for Fps1 56,76.

## Acknowledgements

Dr Adam Wollman, University of Newcastle, UK, for his assistance with the Slimfield data analysis.

## Supplementary information

**Supplementary Table 1.** PCR program settings for mGFP-His3 fragment amplification (tagging) and insertion confirmation (control)

| Stage | Tagging | Control | N of cycles |
| --- | --- | --- | --- |
| Initial denaturation | 98°C, 30 s |  | 1 |
| denaturation | 98°C, 10 s |  | 35 |
| Annealing | 56,5°C, 30 s | 66,3°C, 30 s |  |
| extension | 72°C, 40 s | 72°C, 60 s |  |
| Final elongation | 72°C, 5 min |  | 1 |
| Final hold | 4°C, hold |  |  |

**Supplementary Table 2.**
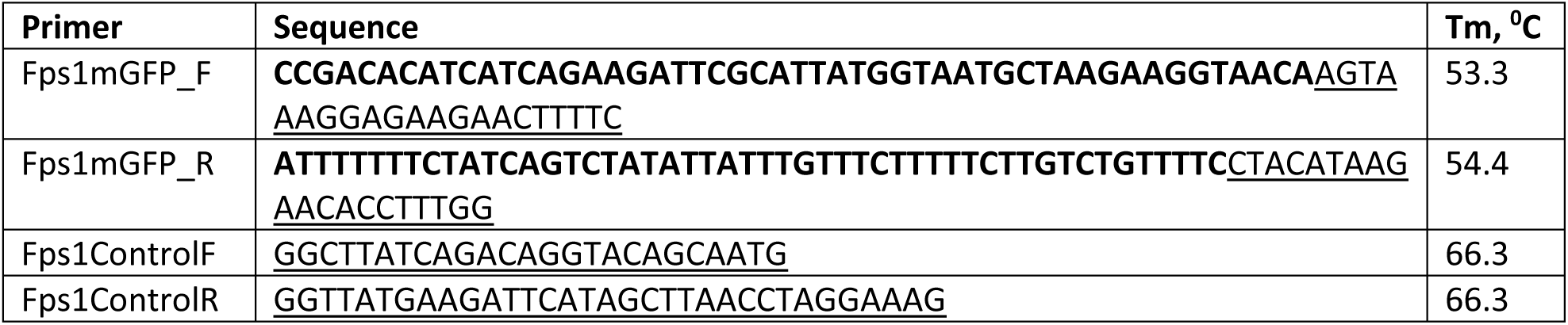
Primers used to amplify mGFP-His3 fragment for further creation of the Fps1-mGFP strain. Noted melting temperatures (Tm) are for underlined sequences only, sequences in bold are 50 bp overhangs to facilitate homologous recombination with *FPS1*. Tms calculated with Thermo Scientifics T_m_ calculator.

**Supplementary Video 1.** Fps1-mGFP foci in living yeast cells under non-stressed conditions.

**Supplementary Video 2.** Fps1.mGFP foci in living yeast cells under hyper-osmotic conditions.

**Supplementary Video 3.** Fps1-mGFP foci in living yeast cells upon oxidative stress.

