## Supplementary figures and images for "Correlating single-molecule characteristics of the yeast aquaglyceroporin Fps1 with environmental perturbations directly in living cells"

### Supplementary Video 1

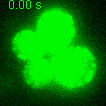

### Supplementary Video 2

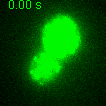

### Supplementary Video 3

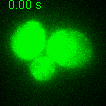
